# A biosafety level 2 surrogate for studying SARS-CoV-2 survival in food processing environmental biofilms

**DOI:** 10.1101/2021.10.29.466519

**Authors:** Austin B. Featherstone, Sapna Chitlapilly Dass

## Abstract

Meat processing plants have been at the center of the SARS-CoV-2 pandemic. There are several factors that contribute to the persistence of SARS-CoV-2 in meat processing plants and one of the factors is the formation of a multi-species biofilm with virus. Biofilm can act as a reservoir in protecting, harboring, and dispersing SARS-CoV-2 from biofilm to the meat processing facility environment. We used Murine Hepatitis Virus (MHV) as a surrogate for SARS-CoV-2 virus and meat processing facility drain samples to develop mixed-species biofilms on commonly found materials in processing facilities (Stainless-Steel (SS), PVC and tiles). The results showed that MHV was able to integrate into the environmental biofilm and survived for a period of 5 days at 7°C. There was no significate difference between the viral-environmental biofilm biovolumes developed on different materials SS, PVC, and tiles. There was a 2-fold increase in the virus-environmental biofilm biovolume when compared to environmental biofilm by itself. These results indicate a complex virus-environmental biofilm interaction which is providing enhanced protection for the survival of viral particles with the environmental biofilm community.

## Introduction

Murine hepatitis virus (MHV) and severe acute respiratory syndrome coronavirus 2 (SARS-CoV-2) belongs to the genus β coronaviruses [1]. Studies suggests that MHV is a good persistent surrogate model for SARS-CoV-2 [1, 2]. In a recent study on wastewater, there were no statistically significant difference between RNA decay of SARS-CoV-2 and MHV[1].

Biofilms in food processing facilities are a major threat to food safety, one of the main carriers of food borne pathogens [3, 4]. Biofilms are multicellular assemblage of prokaryotic and eukaryotic cells that are enclosed in a polysaccharide material and attached to a surface [5]. Bacterial and fungal biofilms have so far been the focus of research on biofilm in food processing facilities, research on the presence of viral particles in the mixed-species biofilm community is spars. There are several factors that could potentially contribute to consider biofilms as an ideal site to harbor SARS-CoV-2 in meat processing facilities. The temperature of the meat processing facilities is maintained at 4-7 °C. The SARS-CoV-2 virions are stable at colder temperatures and stable for several days on stainless steel (SS), copper, plastic, PVC and cardboard [6], which are commonly used materials in meat processing facilities. Thus, making these facilities high risk of harboring and transmitting the SARS-CoV-2 [7]. Although bacteria do not support virus infection, they can promote viral fitness [8]. Specifically, some viruses use components of the bacterial envelope to enhance their stability [8-10]. Moreover, bacterial communities and biofilms can impact the infection of mammals by viruses [8, 9, 11]. Furthermore, from a biophysics perspective, the virus stability could also be enhanced by the thin liquid film produced by bacterial biofilm [6, 12].

There is a critical gap of knowledge in understanding the stability and infectious state of the virus in multi-species biofilms, particularly in meat processing plants. In this study, MHV (SARS-CoV-2 surrogate) was inoculated into meat processing environmental biofilm and developed on SS, PVC, and tiles at 7°C. The MHV survival was analyzed by qPCR and plaque assays to assess the survival rates of the viral particles in environmental biofilms.

## Results

### Biofilm development by MHV in mixed species biofilm from meat processing drain biofilm

Drain sample containing environmental microorganisms was developed on SS (Fig. 1A), PVC (Fig. 1B), and tile surface (Fig. 1C) for five days at 7°C with/without the presence of MHV. The overall mean biofilm microbial cells recovered from the SS chip without MHV ranged from 1.6 to 4.0e+6 CFUs/mL and from the SS chip with MHV ranged from 3.5 to 6.0e+6 CFUs/mL (Fig. 1A). There was a 1.74-fold increase in the biofilm cells developed with MHV on SS when compared to biofilm without MHV. The overall mean biofilm microbial cells recovered from the PVC chips without MHV ranged from 1.8 to 4.0e+6 CFUs/mL whereas the total biofilm microorganisms recovered from the PVC chip with MHV ranged from 5.0 to 7.8e+6 CFUs/mL (Fig. 1B). There was a 2.1-fold increase in the biofilm cells developed with MHV on PVC when compared to biofilm without MHV. The overall mean biofilm microbial cells recovered from the tile chip without MHV ranged from 1.8 to 3.8e+6 CFUs/mL and microorganisms recovered from the tile chip with MHV ranged from 4.8 to 7.0e+6 CFUs/mL (Fig. 1C). There was a 2.11-fold increase in the biofilm cells developed with MHV on SS when compared to biofilm without MHV.

**Fig. 1.**
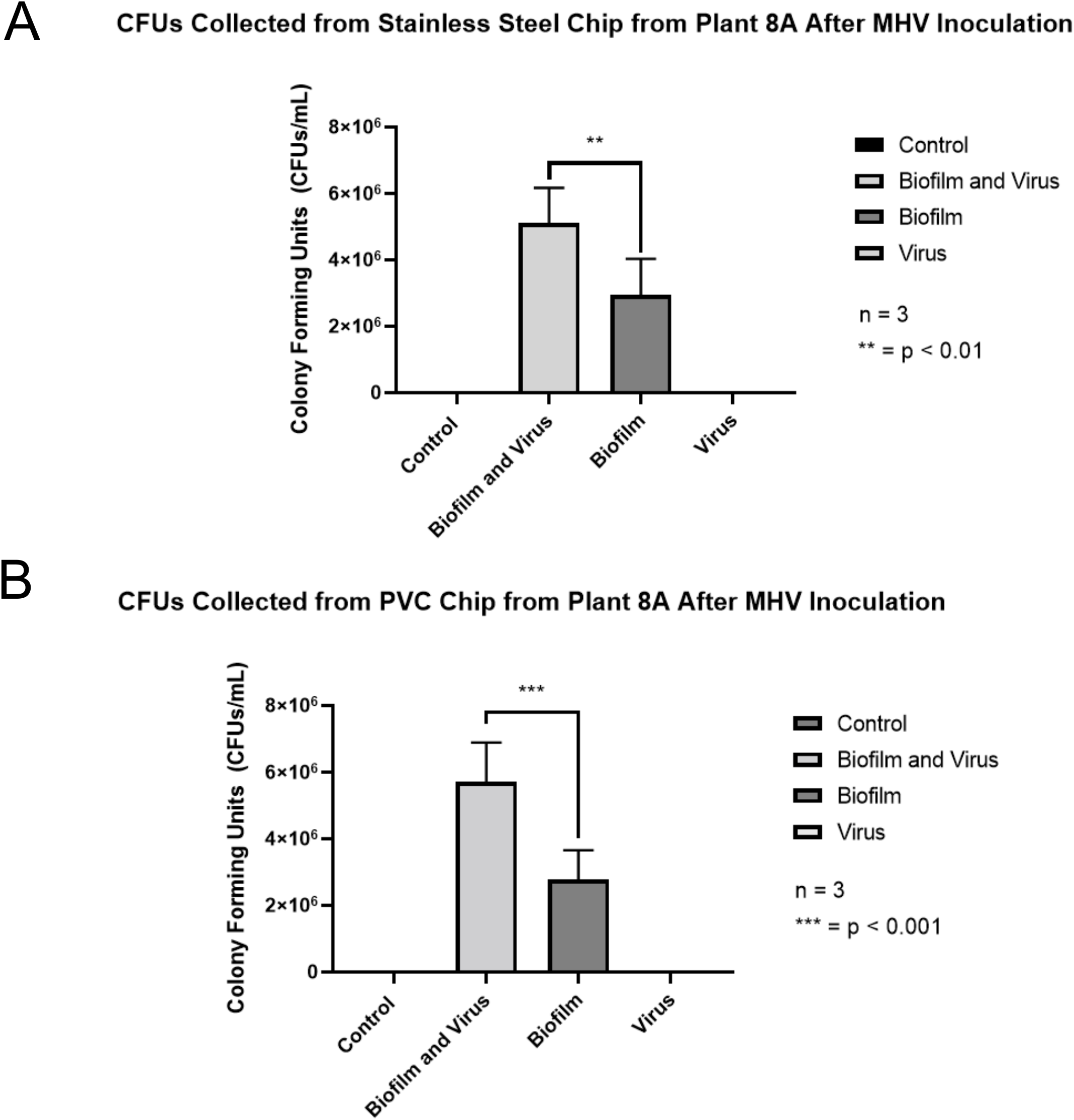
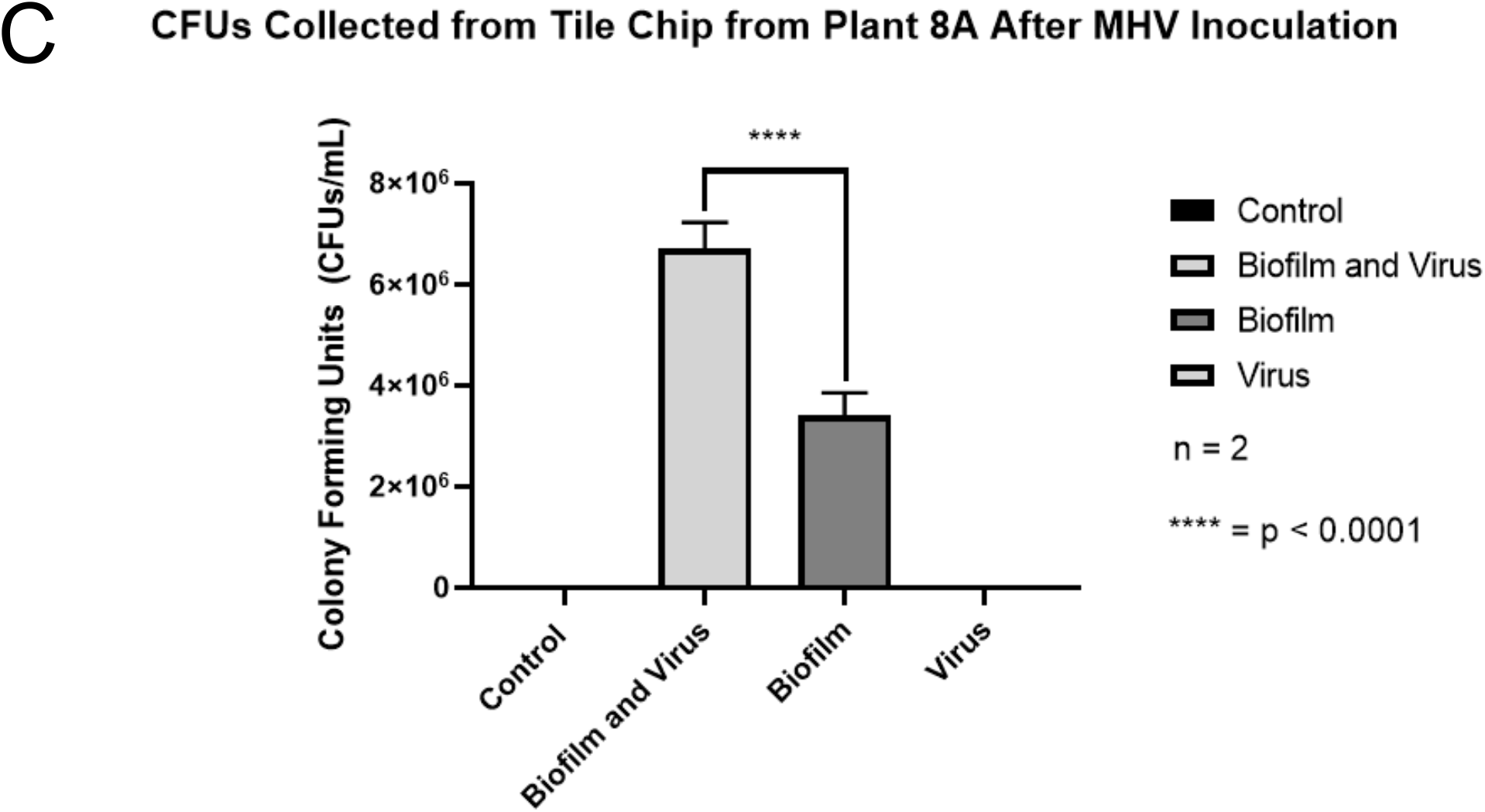
CFU counts from biofilm + virus and biofilm samples on stainless steel, PVC, and tile chips. (A-C) CFU counts for biofilm + MHV and biofilm samples on (A) stainless steel, (B) PVC, and (C) tile chips. Each sample was plated in duplicate. Results in this figure are the mean values and standard deviations (error bars) from three independent experiments. Statistical significance was analyzed by unpaired t-test. n.: not significant; * : P < 0.05; * * * : P < 0.001; * * * * : P < 0.0001.

### qPCR analysis to determine the survival rate of MHV in mixed species biofilm from meat processing drain biofilm

The average critical threshold (CT) for the biofilm and MHV sample on SS chips was 15.4 whereas the CT for virus by itself on SS was 14.2 (Fig. 2A). The difference between the two values was not significantly different by unpaired T-test. The average critical threshold (CT) for the biofilm and MHV sample on PVC chips was 15.6 whereas the CT for virus by itself on PVC was 14.0 (Fig.2B). The difference between the two values was not significantly different by unpaired T-test. The average critical threshold (CT) for the biofilm and MHV sample on tile chips was 22.0 whereas the CT for MHV by itself on stainless steel was 21.6. The difference between the two values was not significantly different by unpaired T-test.

**Fig. 2.**
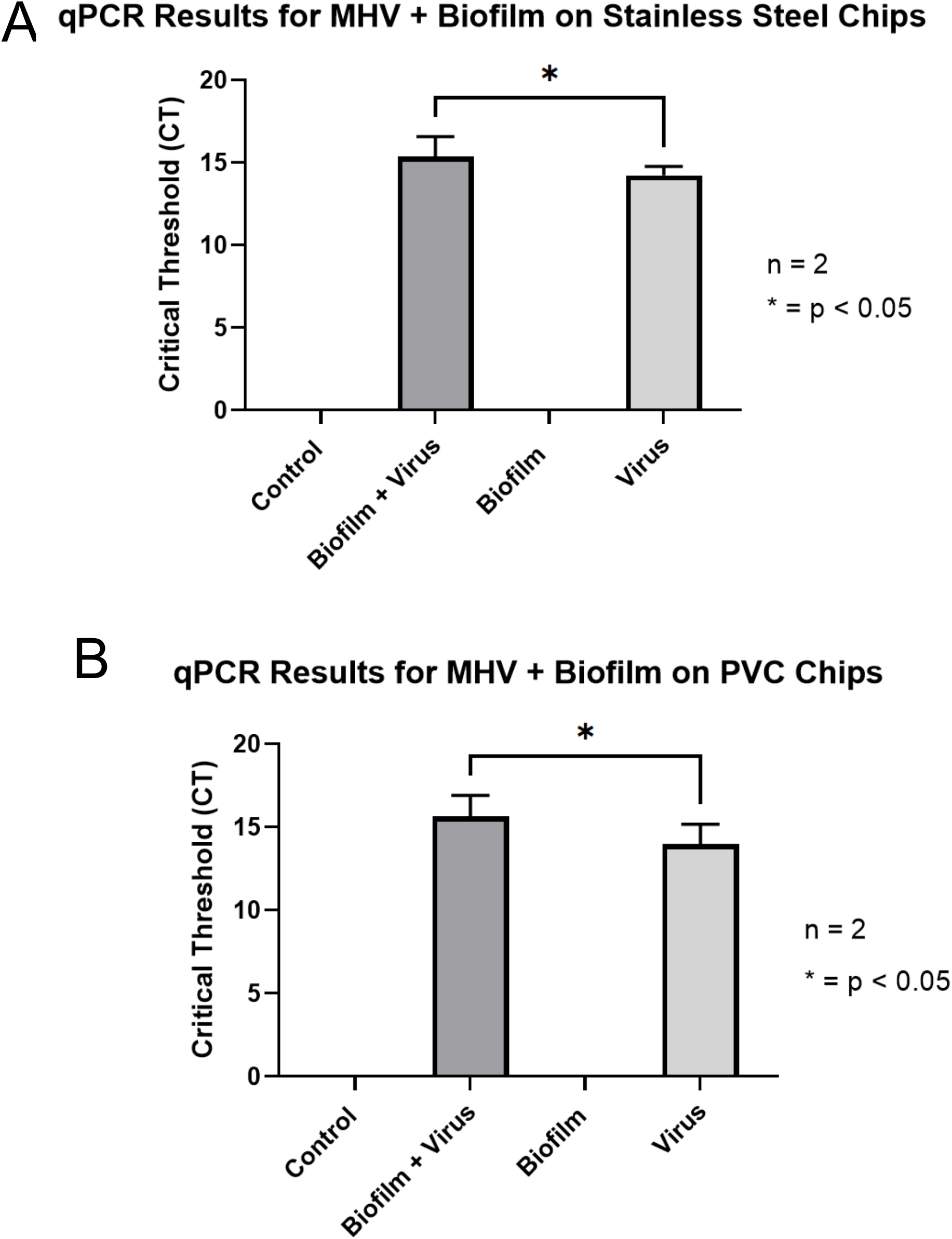
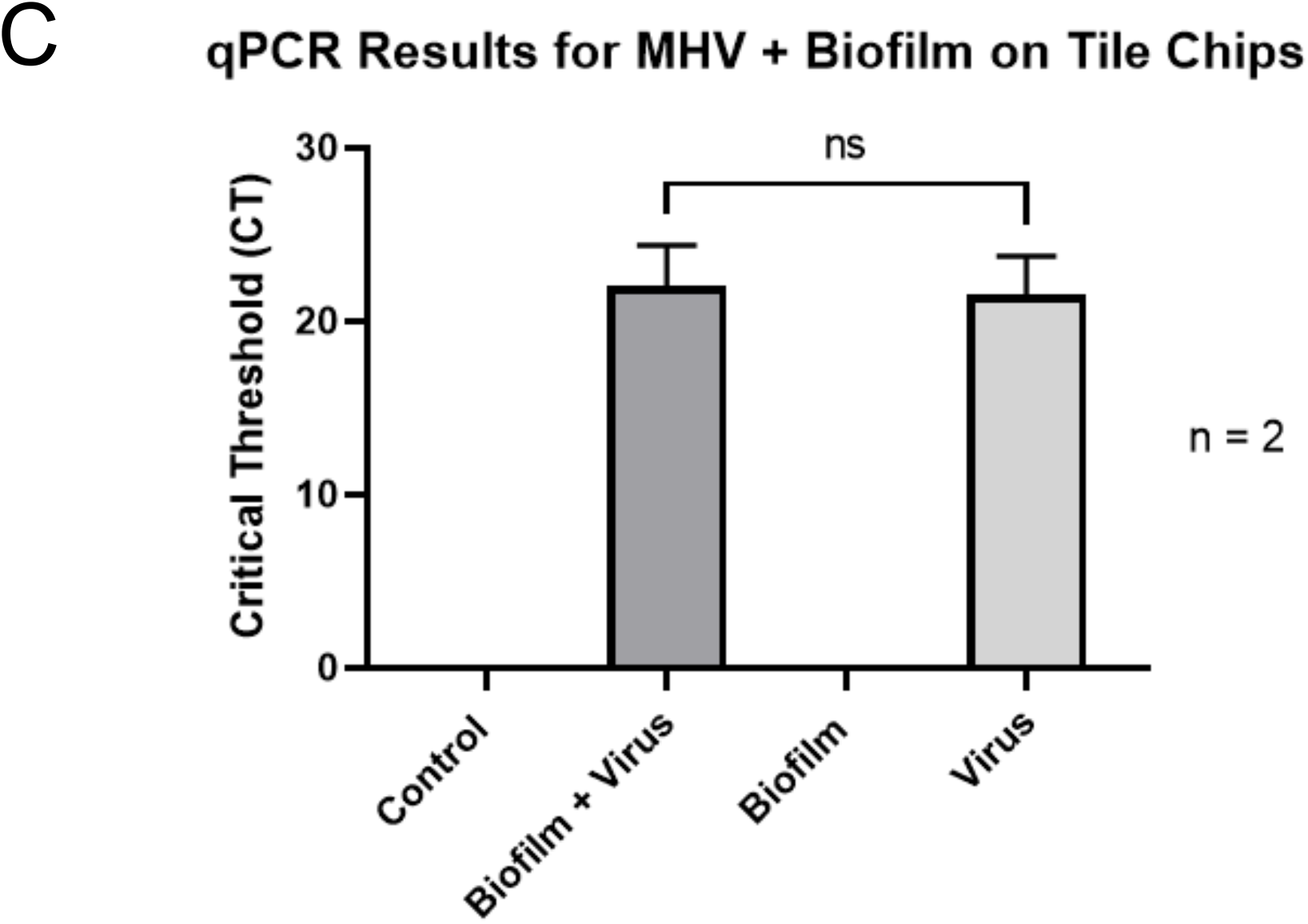
qPCR analysis of MHV mixed with biofilm and pre-incubated for 5 days on stainless steel, PVC, and tile chips. (A-C) qPCR analysis of MHV mixed with environmental biofilm on stainless steel, PVC and on tile chips. 1 × 10^4^ VPUs of MHV were added to a stainless steel chip along with an floor drain biofilm sample collected from the cooler of meat processing plant A. qPCR analysis had the samples analyzed in quadruplicate. Results in this figure are the mean values and standard deviations (error bars) from two independent experiments. Statistical significance was analyzed by unpaired t-test. n.s.: not significant; * : P < 0.05.

### Plaque assay to determine the infectivity of MHV in mixed species biofilm from meat processing drain

The average viral particle units (VPUs)/mL for the biofilm and MHV samples on the SS chips was 650 VPUs/mL, whereas the average VPUs/mL for the MHV sample by itself on the SS chips was 4250 VPUs/mL (Fig. 3A). There was a 6.54-fold difference between the biofilm and MHV VPUs/mL compared to the virus by itself on SS chips. The average VPUs/mL for the biofilm MHV samples on the PVC chips was 600 VPUs/mL, whereas the average VPUs/mL for the MHV sample by itself on the PVC chips was 5500 VPUs/mL (Fig 3B). There was a 9.17-fold difference between the biofilm and MHV VPUs/mL compared to the MHV by itself on PVC chips. The average VPUs/mL for the biofilm and MHV samples on the tile chips was 675 VPUs/mL, whereas the average VPUs/mL for the MHV sample by itself on the tile chips was 6250 VPUs/mL (Fig 3B). There was a 9.26-fold difference between the biofilm and MHV VPUs/mL compared to the MHV by itself on PVC chips.

**Fig. 3.**
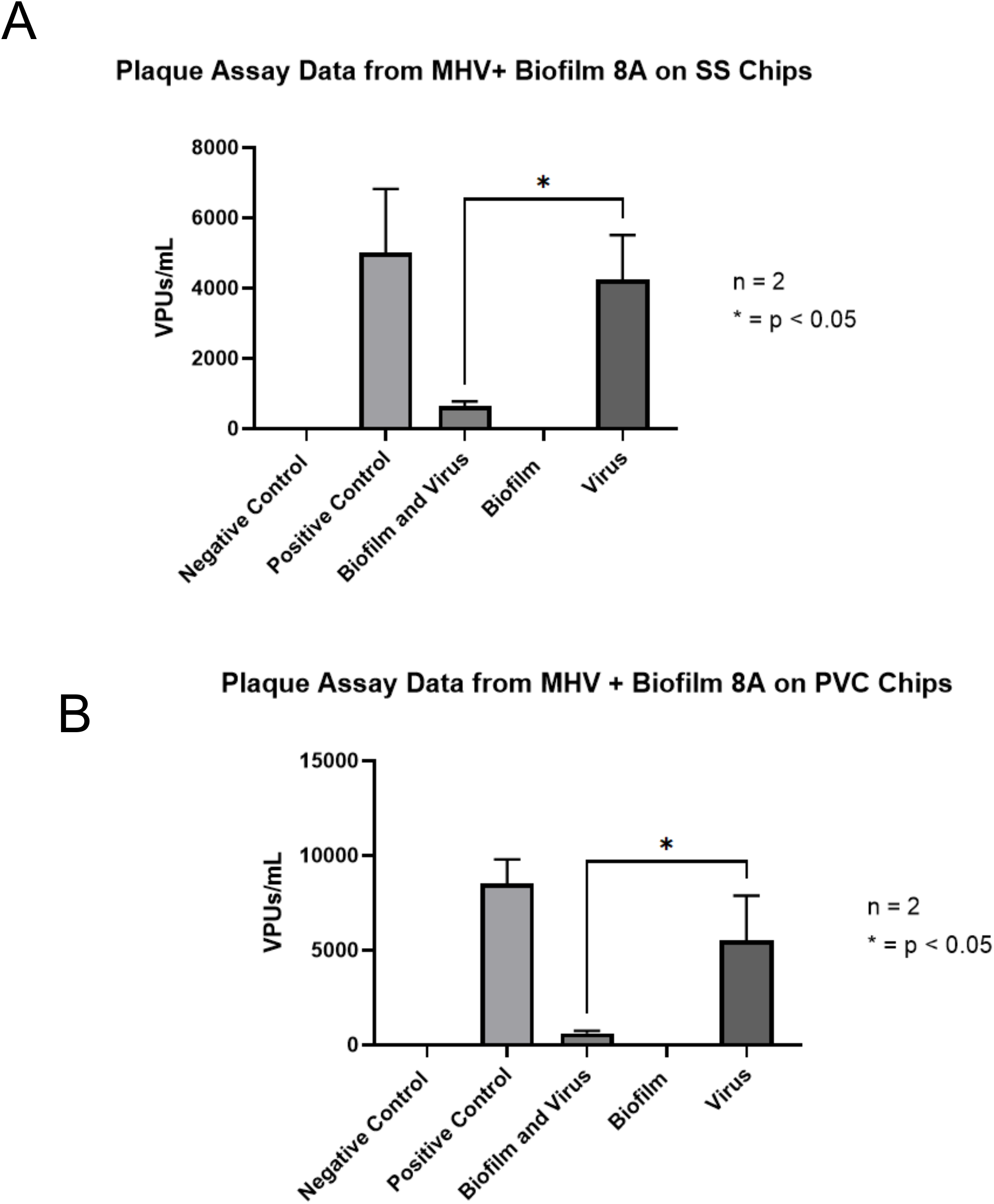
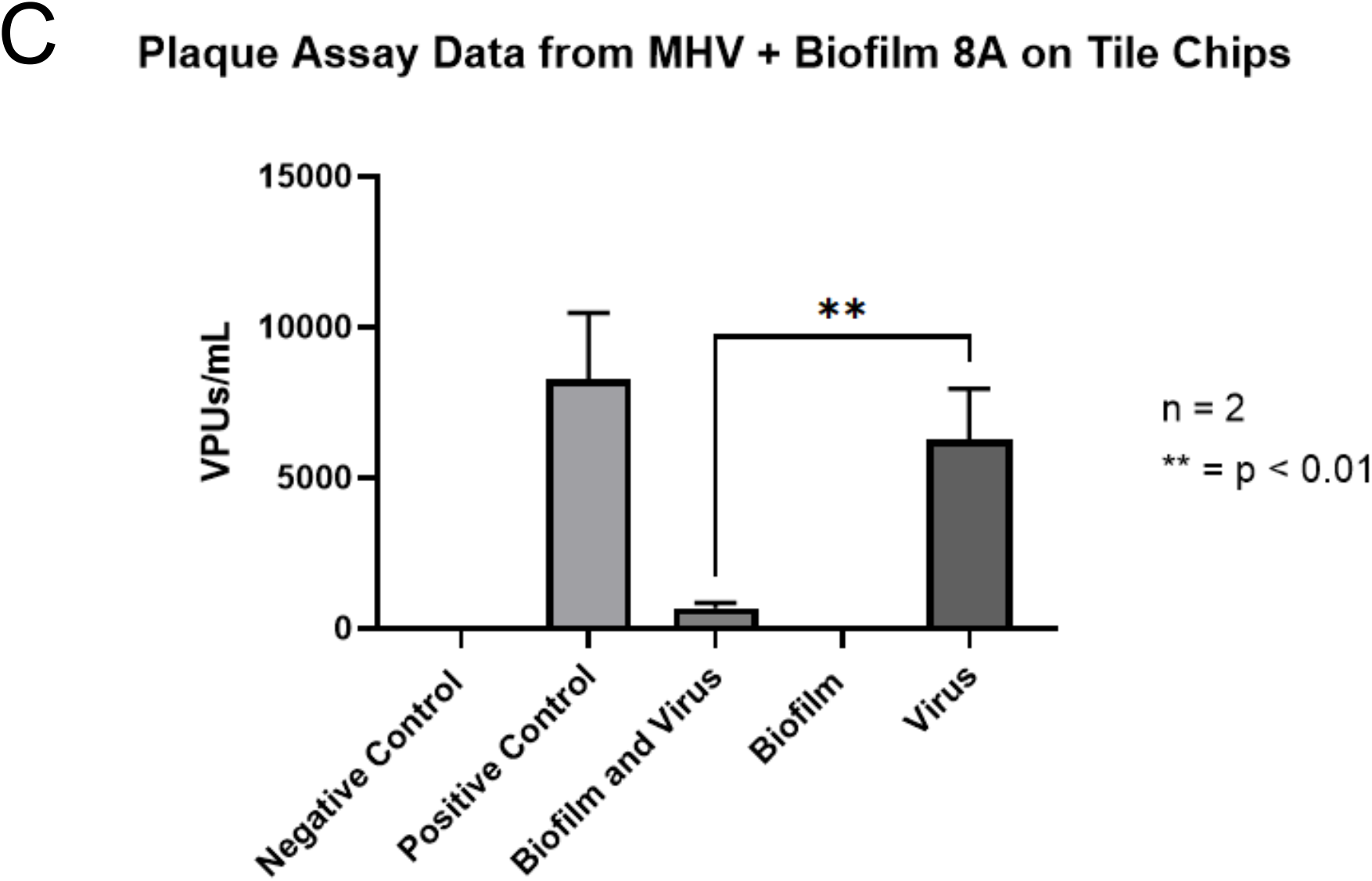
Plaque assay results from biofilm + virus and virus samples on stainless steel, PVC, and tile chips. (A-C) Results from plaque assays on samples collected from (A) stainless steel, (B) PVC, and (C) tile chips. Each sample was plated in duplicate. Results in this figure are the mean values and standard deviations (error bars) from two independent experiments. Statistical significance was analyzed by unpaired t-test. n.: not significant; * : P < 0.05.

## Discussion

The National Cattlemen’s Beef Association [13](predicts that the cattle industry will be potentially experience a $ 13.6 billion loss due to the novel coronavirus SARS-CoV-2, impacting ranchers across the country [13]. This loss stems from the rampant spreading of SARS-CoV-2 virus among meat processing plant workers and resulting in the closure of many meats processing and packing plants. The bottleneck in the supply chain between the livestock producers, feedlot operators and the processors are creating an impending breakdown in the nation’s meat supply. Additionally, the factors that contributes to the persistent high level of COVID-19 infection within the meat processing plant could be the conducive environment for the infectious virus particle to be stable for a prolonged period. The temperature of the meat processing chain are maintained at 4-7°C. The SARS-CoV-2 virions are stable at colder temperatures; hence these facilities are at high risk of harboring and transmitting the SARS-CoV-2. These virus particle can be stable for several days on stainless steel, copper, plastic, PVC and cardboard [6], which are commonly used materials throughout the farm-to-plate chain [7], thus making the meat processing facility a hotspot for SARS-CoV-2 virus.

We utilized MHV as a surrogate for SARS-CoV-2 as they are useful for assessing methods and performance under biosafety level 2 environments [1]. The floor drain samples from meat processing samples were used as a representation of the microbial community representative for meat processing plant. As floor drains collect all rinsing water and liquid wastes in the plants, the microbial communities in the floor drains represent the microecological niches that encompass the various microorganisms in the processing plant environment [14, 15]. We investigated the survival rates of MHV (SARS-CoV-2 surrogate) in meat processing environmental biofilm developed on commonly used materials in the meat processing facilities SS, PVC, and tiles. We observed that the viral RNA integrated with the environmental biofilm on all the materials tested (SS, PVC and tiles) and survived the 5-day incubation at meat processing environment (7°C) temperature. There are several environmental parameters that could have facilitated the survival of MHV in the viral-environmental biofilm. The temperature at the meat processing facility is at 4-7 °C, these temperatures are ideal for the virions to be stable [6]. The survival of the viral RNA was not significantly different when integrated in the environmental biofilm developed on meat processing materials or when it was inoculated on the meat processing materials by itself with viral support media (DMEM). In nature, microbial communities survives as mixed-species communities with complex interactions with eukaryotes and prokaryotes [16] living in complex biofilm web [4, 17, 18]. Therefore, this finding is critical in deciphering the role of biofilm in harboring the viral particles. Given the impact of SARS-CoV-2 on meat processing facilities, it is vital to understand the role of biofilm in serving as the hotspot for viral harborage and dispersion. In addition, we also found that the biofilms biovolumes increased with the addition of the virus to the environmental biofilm when compared to the biofilm by itself. The increase in biovolume could provide enhanced protection to virus from sanitizers. This result is similar to the finding from other work on biofilm where the viral particles enhanced the biovolumes of the biofilm[2, 19-21].

Biofilms are formed in many areas of food processing environments, including floors, drains, difficult to clean surfaces such as the back of the conveyor belts and pipes [22, 23]. These surfaces become hot spots that attract biofilm development due to poor accessibility and difficulty for regular maintenance of hygiene and sanitation maintenance [24]. Furthermore, nearly all biofilms in food processing environments consist of multiple species of microorganisms, and the complex interactions within the community significantly influence the architecture, activity, and sanitizer tolerance of the biofilm [14, 15, 23, 25]. If viral particles present in a meat processing plant are protected within the the biofilms, regular sanitizer treatment will not eliminate the biofilm or virus[4, 14, 26, 27].

In addition, to providing sheltered and protected living for viral particles from sanitizers, microbial communities in the biofilm could also enhance the dispersal of viruses through several active processes: Viruses can bind to bacteria [9], and they can be transported over large distances by bacterial motility. The bacteria can also colonize new areas by continuously producing new extracellular matrix (ECM), actively swarming outwards [14, 15], thus creating new locations where the virus can harbor throughout the meat-processing facility. Even without binding to the viruses, bacteria will generate strong flows [16] that mix the surrounding fluid [17]. Therefore, viral particles could be spread rapidly from biofilms in the meat facility. Microbial biofilms should thus be considered as potential reservoirs of pathogenic viruses. Indeed, they are probably responsible for numerous persistent viral infections [20]. Thus, biofilms should thus be considered as potential reservoirs of pathogenic viruses. SARS-CoV-2 is well documented to be airborne and spread through aerosols [6, 28]. Hence, once the virus is airborne, the colder temperature and the airflow from HVAC systems in the meat processing facility will enable the spread of the SARS-CoV-2 throughout the plant, in agreement with recent fluid mechanics simulations [29].

The above findings led us to conclude that multi-species biofilms may provide a safe reservoir for SARS-CoV-2 to persist as an infectious agent due to the advantages conferred by the biofilm structure and facilitate the dispersal of the viral particle through the collective microbial motility. This could be one of the reasons for the higher rates of COVID-19 cases in meat processing facilities. Due to the nature of the work, it will be difficult with workplace physical distancing, personal hygiene, crowded living and transportation conditions among the meat processing frontline workers [30].

## Conclusions

In summary, our results suggest that environmental multi-species biofilm from meat processing plant can harbor viral particles (MHV which is used as a surrogate for SARS-CoV-2) and survive for prolonged periods of time protecting them from sanitizers. Thus, facilitating a safe reservoir for virus to persist and periodically disperse the viral particle to the meat processing facility, thus making these facilities the hotspot for higher COVID 19 infections. Further studies will be required to decipher molecular mechanism of how these interaction takes place, and this knowledge will help in designing targeted intervention strategies to reduce the harborage and spread of viral lodged multi-species biofilms.

## Materials and Methods

### Drain sample collection and characterization

The floor meat processing drain biofilms were collected following the previously described protocol [14]and were generously provided for this study by Drs. Mick Bosilivac and Rong Wang USDA-ARS-USMARC, Clay Center, Nebraska. The biofilm used in this study was collected from floor drain biofilm from the cooler of a meat processing facility Plant A.

### Cell lines and MHV propagation

L2 cells (ATCC® CCL–149TM) were used for MHV plaque assay. In addition, the mouse asterocytoma-derived cell line (DBT), was used to propagate MHV (generously provided by Dr. Julian Leibowitz, Texas A&M Health Science Center, College Station, TX). All cells used in this study were cultured at 37°C in 5% CO_2_ in Dulbecco’s modified Eagle medium (DMEM; Cellgro) supplemented with 10% fetal bovine serum (FBS), penicillin (50 IU/mL), and streptomycin (50 µg/mL). MHV strain A59 (ATCC® VR-764) were used for all experiments. The virus stocks used for infectivity were produced as previously described[31].

### Assay of MHV infectivity

The viral infectivity for each virus was determined by titrating each virus stock onto cultured L2 cells and performed a solid double overlay plaque assay as previously described [32].

### Culture conditions for drain sample and MHV

MHV stocks were cultured to a viral titer of 10^4^ VPUs/mL prior to the start of the experiment [32]. To simulate the meat processing plant environment, floor drain samples were inoculated into the biofilm was 50-fold inoculated into Lennox Broth (LB, Acumedia Manufacturers, Baltimore, MD) without salt (LB-NS) medium and incubated at 7 °C for 5 days with orbital shaking at 200 rpm [14]. On the fifth day, a 1.0 mL aliquot was removed from each sample, diluted in sterile LB-NS medium, and plated onto Trypticase soy agar (TSA) for colony enumeration after overnight incubation at 37 °C.

### Biofilm formation with drain sample and MHV

To investigate whether biofilm formation with floor drain samples can support the harborage of MHV, biofilms by the drain microorganism with or without MHV and MHV alone and control were developed on SS chip, PVC, and floor tiles (Fig 4). The experiments were set-up in duplicates in a 6 well plate. Each well had one sterile SS/PVC, or tiles chips (18×18×2 mm): (A) Drain sample + virus: 100 µL of the 5 day culture (described above) was placed on the SS/PVC or tiles chip along with 100 µL of virus in DMEM and 100 ul of LB-NS media. (B) Drain samples: 100 µL of the 5 day culture (described above) was placed on the SS/PVC or tiles chip. (C) MHV alone: 100 µL of MHV in DMEM (D) Control: 100 ul of DMEM and 100 ul of LB-NS. The biofilm set-up was incubated at 7°C for 5 days.

**Fig. 4.**
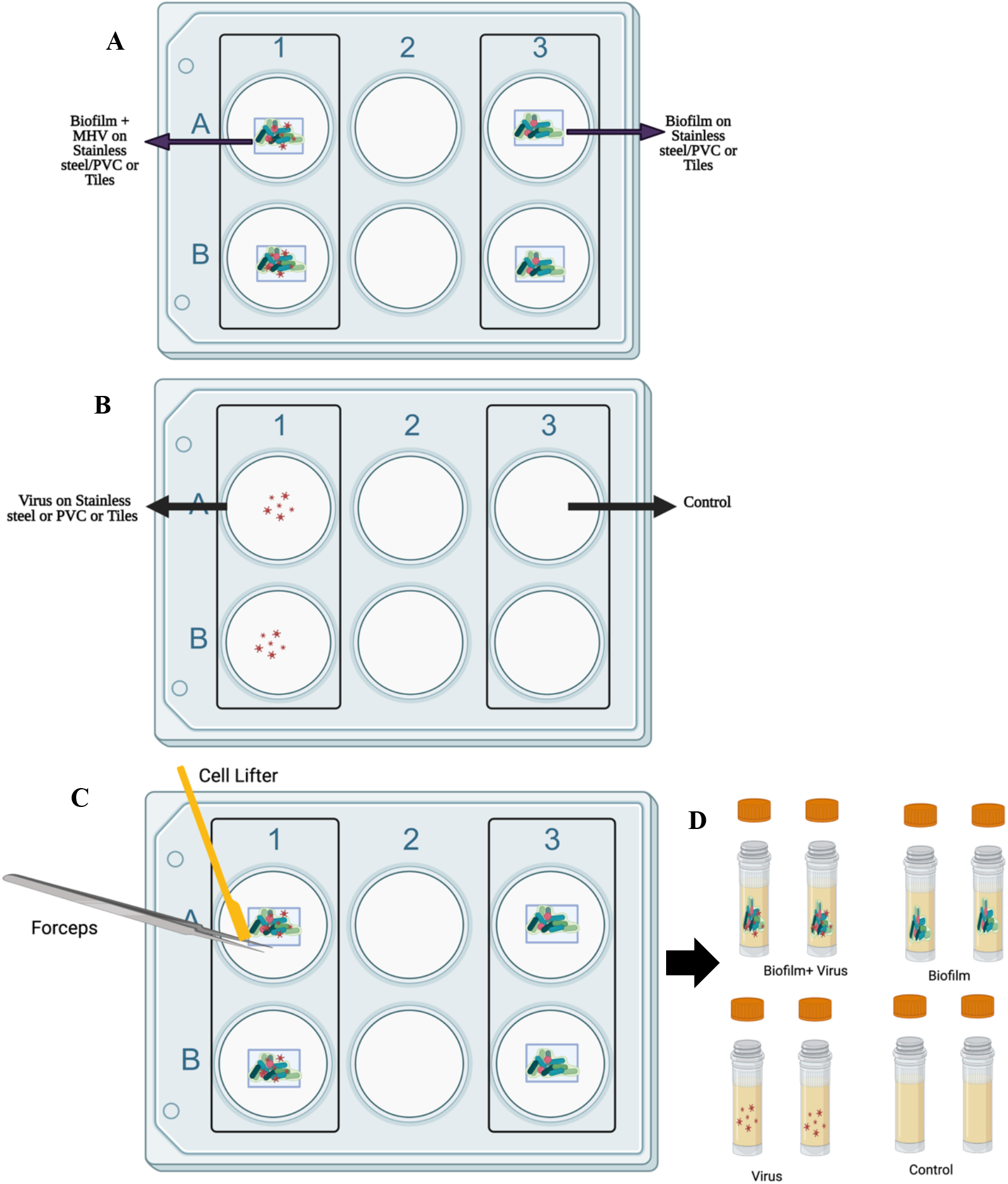
Schematic representation of the biofilm+ MHV inoculated study on SSl/PVC/Tiles. (A and B): Experimental set up with Biofilm+ Virus, Biofilm, Virus and Control in duplicates. The experimental set is incubated at 7 C for 5 days. (C). After 5 Days the biofilm is harvested from SS/PVC/Tiles using a cell lifter and forceps and rinsed with 1000 ul of LB-NS, the support materials after scraping (SS/PVC/Tiles is removed. (D) Harvested cells are stored in a screw-cap tube in −80C until further use.

At the end of the incubation period, each chip was harvested for the biofilm biomass (Fig 4) by lifting the chip with sterile forceps, scraping biofilm from both sides with a sterile cell scraper and rinsing the chip with 1000 µL of LB-NS.

The homogenate was mixed several times by pipetting. The drain biofilm biomass was determined by 10 fold dilution of the homogenate in LB-NS and plated on TSA plates for colony enumeration after overnight incubation at 37C. The remaining homogenate was used for qPCR and plaque assay analysis.

### Viral RNA Extraction

Viral RNA from each samples were extracted and purified to perform RT-qPCR to determine the relative copy numbers for MHV. Viral RNA was extracted using NEB’s Monarch Total RNA Miniprep Kit and following the Tough-to-Lyse procedure following manufacturers protocol. Purified RNA samples were quantified by using a Thermo Fisher Scientific ND-1000 spectrophotometer. RNA samples were stored at −20°C.

### MHV RT-qPCR Analysis

Taqman-based RT-qPCR analysis was completed using NEB’s Luna^®^ Universal Probe One-Step RT-qPCR kit. Purified RNA extracted from MHV virus was used for the positive control and standard curve. The RT-qPCR analyses were completed in 25 µL reactions using the Luna Universal Probe One-Step Reaction Mix. The RT-qPCR mixture contained 10 µL of Luna Universal Probe One-Step Reaction Mix, 1 µL of Luna WarmStart RT Enzyme Mix, 400 nM of forward primer (5’-GGAACTTCTCGTTGGGCATTATACT-3’), 400 nM of reverse primer (5’-ACCACAAGATTATCATTTTCACAACATA-3’), 200 nM of probe (IDT) (5’-FAM-ACATGCTAC-ZEN-GGCTCGTGTAACCGAACTGT-3IABkFQ-3’), 250 ng RNA, and nuclease free water. The RT-qPCR analysis was performed using a Bio-Rad CFX96 Deep Well Real Time thermal cycler. Thermal cycling conditions were set to 55°C for reverse transcription for 10 minutes, denaturation and *Taq* polymerase activation at 95°C for 1 minute, and 40 cycles at 95°C for 15 seconds followed by 60°C for 30 seconds for data collection. RT-qPCR reactions were performed in quadruplicate for each sample and the sample quantification cycle (Cq) was used for data analysis. Data for each sample was compared using positive and negative controls performed in duplicate.

### Plaque assay analysis

Plaque assay from biofilm homogenate were performed to determine the MHV infectivity rate in biofilm. 300 µL of each biofilm homogenate samples were filtered through a .2 µm filter before being serially diluted in DMEM with 2% FBS and 1% Streptomycin/Penicillin mix. Previously published protocol was followed for the double layer overlay plaque assay[32].

## Acknowledgements

The authors would like to thank USDA-NIFA 2020-67015-32330 grant for support of this study. We would like to thank Drs. Mick Bosilivac and Rong Wang (USDA-ARS-UAMRC, Clay Center-Nebraska) for providing meat processing plant biofilms, Dr. Julien Leibowitz (Texas A&M) for providing original stocks of MHV and DBT cell line, and Dr. Amanda Claire Brown (Texas A&M) for help with the initial design of the experiments.

